# Whole-brain optical access in small adult vertebrates with two- and three-photon microscopy

**DOI:** 10.1101/2021.12.09.471956

**Authors:** Najva Akbari, Rose L Tatarsky, Andrew H Bass, Chris Xu

## Abstract

Although optical microscopy has allowed us to study the entire brain in early developmental stages, access to the brains of live, adult vertebrates has been limited. *Danionella*, a genus of miniature, transparent fish closely related to zebrafish has been introduced as a neuroscience model to study the entire adult vertebrate brain. However, the extent of optically accessible depth in these animals has not been quantitatively characterized. Here, we show that two- and three-photon microscopy can be used to access the entire depth of the adult wild type *Danionella dracula* brain without any modifications to the animal other than mechanical stabilization. Three-photon microscopy provides high signal to background ratio and optical sectioning through the deepest part of the brain. While vasculature can be observed with two-photon microscopy, the deeper regions have low contrast. We show that multiphoton microscopy is ideal for readily penetrating the entire adult brain within the geometry of these animals’ head structures and without the need for pigment removal. With multiphoton microscopy enabling optical access to the entire adult brain and a repertoire of methods that allow observation of the larval brain, *Danionella* provides a model system for readily studying the entire brain over the lifetime of a vertebrate.

## Introduction

Understanding the brain requires studying the complex spatiotemporal relationship between neurons and how they come together to give rise to behavior. High scattering and absorption of tissue limits the penetration depth of high-resolution optical imaging inside the brain [1,2]. Small animals such as zebrafish larva and frog tadpoles allow non-invasive, *in vivo* access to vertebrate brains due to relative transparency of the skin and late ossification of the skull [3–11]. However, they often lack the behavioral complexity that develops in adulthood, especially in social contexts related to reproduction such as courtship, copulation, and resource defense. Optical access to adult vertebrate brains with high resolution is currently limited by penetration depth of available techniques and the head size of most commonly-used species [12– 14].

Multiphoton microscopy (MPM) has enabled deep optical access into scattering biological tissues [2,14– 17]. These techniques use longer excitation wavelengths compared to one-photon excitation (e.g. confocal microscopy) which increases the penetration depth of photons and the nonlinear excitation suppresses the out-of-focus fluorescence, improving the contrast. Two-photon microscopy (2PM) has become the gold standard of deep *in vivo* imaging and has greatly advanced our understanding of biological systems [15,18]. Three-photon microscopy (3PM) enables unprecedented imaging depths including through the intact skull of adult mice and zebrafish [2,19–21]. The imaging depth of MPM is ultimately limited by tissue scattering and absorption, therefore, deep access to the brains of larger animals such as mice is limited [12–14].

Smaller adult vertebrates with translucent young including zebrafish (*Danio rerio*) and the smaller relatives of zebrafish, *Danionella*, offer the prospect of imaging throughout the brain of a vertebrate as well as throughout life [19,20,22,23]. *Danionella dracula* adults, studied here, reach approximately 11 to 18 mm in length and have a transparent body with a poorly ossified skull above the brain [24,25]. Like other *Danionella* species, they produce sounds during social interactions making them especially promising for identifying neural mechanisms underlying adult vertebrate reproductive behavior [25].

Recently, optical access to the adult brain of zebrafish (*Danio rerio*) and multiple *Danionella* species (closely related to zebrafish), has been demonstrated with MPM [19,20,22,23]. 3PM allows penetration through the skull and scales of the adult zebrafish head, enabling one to address a new range of questions in this well-established model animal. For example, this permitted structural and functional imaging through the telencephalon and deeper into the optic tectum and cerebellum than was possible before [20]. However, imaging the adult zebrafish brain in its deepest regions has not been achieved yet. While imaging through the *D. dracula* brain to visualize vasculature was also demonstrated with 3PM, the performance of this technique was not characterized [20]. Additionally, optical imaging into the brain with 2PM, which is currently a more widely used technique, has remained relatively shallow.

Here, we demonstrate optical access to the deepest regions in adult *D. dracula* brain with both 2PM and 3PM. We compare 2PM at 920 nm to 3PM at 1280 nm excitation wavelength by imaging fluorescein-labeled vasculature through the deepest part of the adult brain. We additionally compare 2PM and 3PM images obtained with the same excitation beam at 1280 nm by simultaneous labeling of the blood vessels with fluorescein and Alexa Fluor 680. Our results indicate that the higher order of excitation in 3PM is greatly beneficial for maintaining high contrast and optical sectioning at greater depths than with 2PM. Moreover, we show that 2PM and 3PM are capable of readily penetrating the entire brain of this wild-type, pigmented vertebrate without the need for any modifications to the animal other than mechanical stabilization.

## Results

### Blood vessels are resolvable throughout the deepest part of the brain

We labeled the vasculature of adult *D. dracula* with dextran-coupled fluorescein and collected 2PM and 3PM images using 920 nm and 1280 nm excitation wavelengths, respectively (see Methods). Using the white-light path of the microscope (Fig. 3C), we chose an imaging region at the boundary of the cerebellum and optic tectum to ensure that the deepest part of the brain (hypothalamus) was within the field-of-view (FOV). To determine the bottom of the brain, we collected the second harmonic generation (SHG) signal generated by the 1280 nm excitation wavelength. A prior anatomical study enabled us to clearly identify the bone at the bottom of the brain in the SHG channel (Fig. 4) [24]. To compare the performance of 2PM and 3PM, we collected images through the entire depth of the brain. The powers used to obtain the deepest images were 14 mW and 16 mW for 2PM and 3PM, respectively, for similar pixel values (within ∼25%). 3PM maintained high signal-to-background ratio (SBR) throughout the entire depth of the brain, up to 945 µm at the cerebellar-tectal boundary (Fig. 1A). In comparison, 2PM images had substantially lower SBR in the deep regions of the brain (Fig. 1A,B).

**Figure 1:**
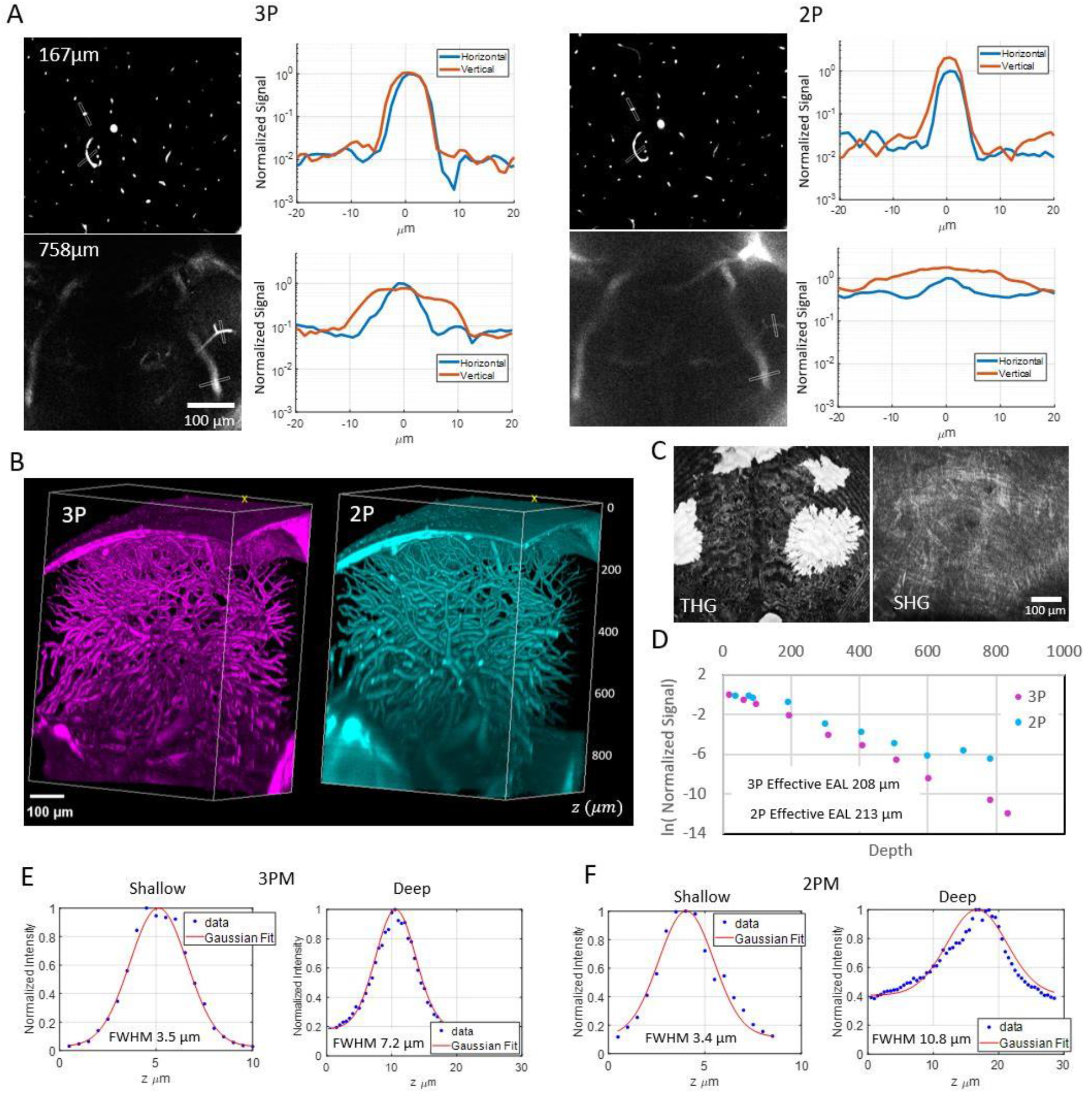
Characterization of 2PM and 3PM images through the deepest part (midbrain) of the adult *D. dracula* brain. Excitation wavelengths of 920 nm at 2 MHz repetition rate and 1280 nm at 333 kHz repetition rate were used for 2PM and 3PM, respectively. A) Signal to background ratio comparison of horizontal and vertical blood vessels for 2PM and 3PM images at two depths inside the brain. In each line profile plot, the values are normalized to the maximum brightness of the horizontal blood vessel. B) Representative 3D reconstruction of 3PM (left) and 2PM (right) images of fluorescein-labeled blood vessels in the midbrain. C) Maximum projection of third harmonic generation (THG, left) and second harmonic generation (SHG, right) of all frames containing the skin. Signals were generated by 1280 nm excitation. Pigments produce very bright signal in the THG channel. D) Characterization of effective attenuation length inside the brain for 2PM and 3PM excitation wavelengths as described in the methods section. E) Axial width of small blood vessel characterized in shallow (∼100 µm) and deep (∼600 µm) regions with 3PM. F) Axial width of small blood vessel characterized in shallow (∼100 µm) and deep (∼600 µm) regions with 2PM.

We measured the lateral and axial width of a small horizontal blood vessel in shallow (∼100 µm) and deep (∼600 µm) regions. In shallow regions, we found the lateral width to be 2.9 µm for both 3PM and 2PM and the axial width to be 3.4 µm and 3.5 µm for 3PM and 2PM, respectively. In the deep regions, we found the lateral width to be 2.7 µm and 4 µm and the axial width to be 7.2 µm and 10.8 µm for 3PM and 2PM, respectively (Fig. 1E,F). Additionally, we compared the brightness of horizontal blood vessels to vertical blood vessels for both 3PM and 2PM (Fig. 1A). While 3PM images maintain equal brightness for horizontal and vertical blood vessels, in deep 2PM images, the brightness of vertical blood vessels was higher than that of horizontal ones. The increased brightness in vertical blood vessels when compared to the horizontal ones in the 2P images can be attributed to distortions in the point spread function (PSF), causing elongation of the focal volume in the axial direction and therefore increasing the excitation volume [21]. With NA=1 as in our system the fluorescence generated within a 5 µm thick slab around the focal plane is ∼89% of the total fluorescence from an infinite volume for 2P excitation (920 nm) and ∼99% for 3P excitation (1280 nm). Therefore, the fluorescence generated by vertical blood vessels is expected to be approximately 1.1 times that generated by the horizontal ones in 2PM images. However, the brightness in vertical blood vessels was measured to be ∼2 times the brightness of horizontal ones for 2P excitation, indicating degraded axial confinement of 2PM. This observation is further supported by the effective attenuation length (EAL) values from these images which are measured to be 208 µm and 213 µm for 1280 nm and 920 nm excitation wavelengths, respectively [21].

While most of our experiments were done with low repetition rate lasers, we also used an 80 MHz laser for 2PM excitation to evaluate the power levels necessary for imaging through the brain. We collected 2PM and 3PM images of three adult fish through the deepest part of the brain using 80 MHz repetition rate for 2PM excitation. Differences in horizontal and vertical blood vessel brightness were observed, indicating PSF degradation (Supp. Fig. 2). We measured the EAL (see Methods) to be ∼ 267 µm at 920 nm and ∼ 300 µm at 1280 nm. To image (with similar pixel brightness values) the deepest part of the brain average power levels of 213 mW and 18 mW were used for 2PM and 3PM, respectively. Fish had healthy body color and movement immediately after imaging and for at least one week after imaging.

### 3PM improves image quality by suppressing the side lobes of the PSF

To delineate the effects of a longer wavelength and the higher order nonlinear excitation, we compared 2PM and 3PM both with 1280 nm excitation by injecting blood vessels with a combination of Alexa Fluor 680 and Fluorescein (both dextran coupled). Maximum powers used to obtain images through the deepest part of the brain were 1.2 mW and 34 mW for 2PM and 3PM, respectively, with pixel values about 50% higher in 2PM images.

In deep 2PM images, the brightness difference between horizontal and vertical blood vessels was observed (Fig. 2A) indicating degradation of the PSF. This effect is also observable as blurriness of the features in the deep regions of the maximum projection of the 2PM stack but not in 3PM (Fig. 2C). The EAL value obtained from 2PM images was larger than the value obtained by analyzing the 3PM images, further verifying PSF degradation of 2PM [20] (Fig. 2B). We measured the lateral and axial width of a small horizontal blood vessel in shallow (∼100 µm) and deep (∼690 µm) regions by collecting the 2PM and 3PM signals from the same blood vessel (Supp. Fig. 3). In shallow regions, we found the lateral width to be 2.6 µm and 3.3 µm and the axial width to be 4.2 µm and 6.5 µm for 3PM and 2PM, respectively. In deep regions, we found the lateral width to be 5.8 µm and 7.3 µm and the axial width to be 8 µm and 12.1 µm for 3PM and 2PM, respectively (Supp. Fig. 3).

**Figure 2:**
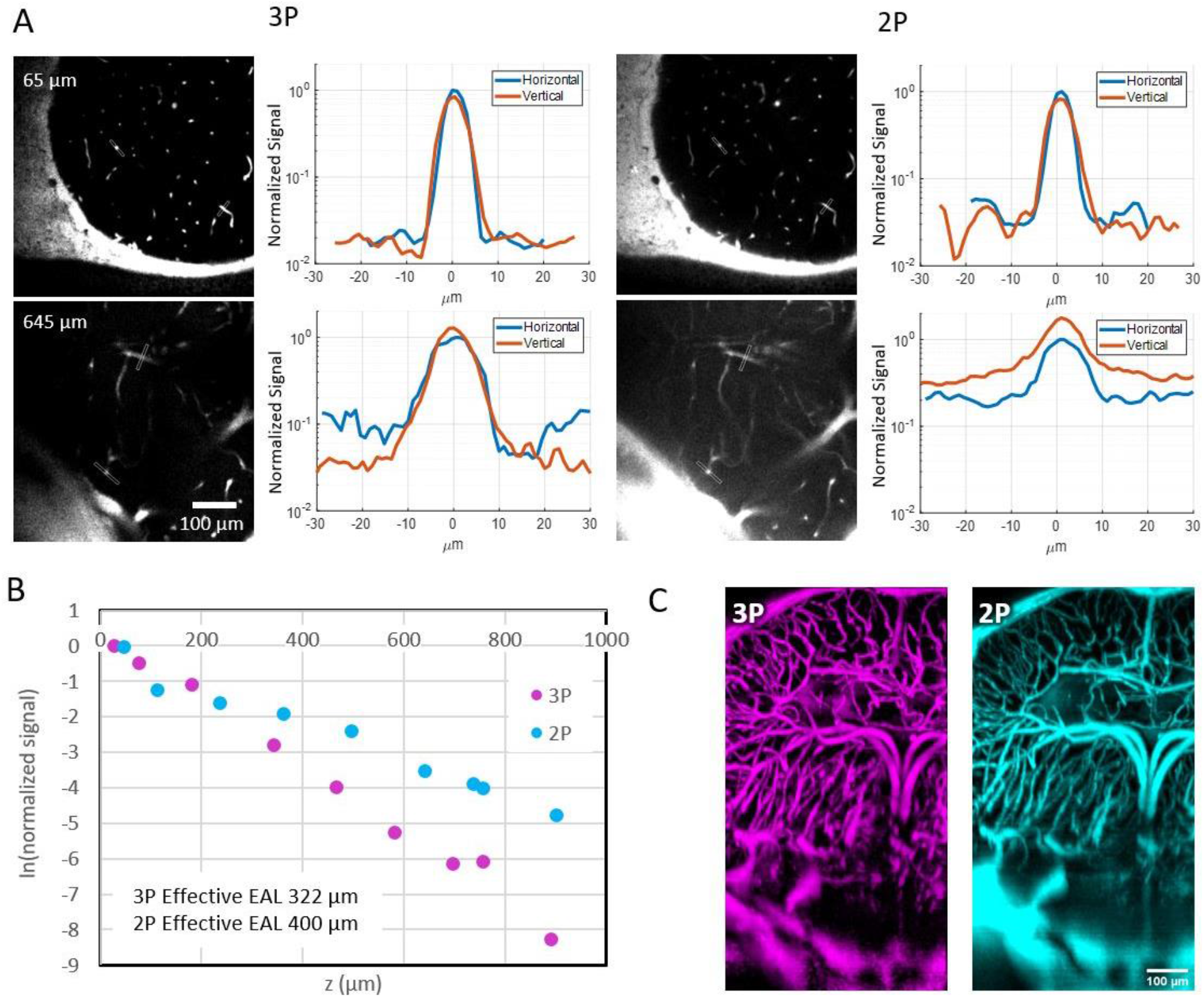
Characterization of 2PM and 3PM (both with 1280 nm excitation) images through the deepest part (midbrain) of the adult *D. dracula* brain. A) Brightness comparison of horizontal and vertical blood vessels for 3PM and 2PM images at various depths inside the brain. In each line profile plot, the values are normalized to the maximum brightness of the horizontal blood vessel. B) Characterization of effective attenuation length inside the brain for 2PM and 3PM excitation wavelengths as described in the methods section. EAL is measured to be 400 µm with 2PM images and 322 µm with 3PM. C) Maximum projection of a column containing the deepest part of the brain.

## Discussion

We demonstrate MPM through the entire brain of an adult vertebrate, *D. dracula*, with high resolution and contrast without the need for any modifications to the animal other than mechanical stabilization. Deep, high-resolution imaging in small animal species like the one studied here allows us to observe an intact adult vertebrate brain throughout its entire depth. While both 2PM and 3PM can penetrate deep inside the brain, 3PM maintains high contrast through the deepest part of the brain. By comparing both 2PM and 3PM with the excitation wavelength of 1280 nm, we show that using longer wavelengths for 2PM is not sufficient for improving image contrast in the deep regions of the brain.

To determine the source of the background in deep images we quantified the volume distribution of blood vessels in different regions of the brain which are the main source of fluorescence in our images. The quantification of blood vessel distribution (Supp. Fig. 4) shows that in all regions of the brain, blood vessels occupy less than ten percent of the volume. Based on the effective attenuation lengths obtained from tissue and the fluorophore distribution in blood vessels (≤10% fluorescent labeling density), we would expect that all experimental depths (Fig. 1 and 2) should have much higher SBR for both 2PM and 3PM in typical imaging conditions [12,13].

In small aquatic vertebrates such as *D. dracula* (and zebrafish), some fluorescence is also generated by pigmentation, particularly from the eyes, that contributes to the background (Supp. Fig. 5 and 6). Although zebrafish genetic lines that remove pigments can help improve contrast, the complete removal of pigmentation can cause health issues (e.g. blindness) and therefore can limit applicability to biological questions [26,27]. While decreasing fluorophore volume fraction (e.g. pigment removal) can decrease the uniformly distributed background (i.e. bulk background [14]), the differences in horizontal and vertical blood vessel brightness cannot be explained by fluorescent labeling density alone.

Aberrations in a sample can distort the PSF, leading to localized brightness differences and therefore, background generation that would depend on the specific location in the field of view (i.e. defocus background [14]). For example, the baseline of the deep 2PM image is different between horizontal and vertical blood vessels in Fig. 2A, indicating a potential localized background contribution. In small animals such as *D. dracula*, tissue inhomogeneities throughout the cone of light can lead to degradation of the PSF, producing defocus background in 2PM images that significantly degrade the SBR. Due to its higher order of excitation, 3PM can maintain a sharper focus through highly aberrating samples. This effect was recently characterized in through-skull imaging of the mouse brain [21]. While scattering and aberration are mainly caused by a single layer of dense bone in through-skull imaging of the mouse brain, aberrations in *Danionella* likely result from multiple tissue types that enter the cone of light with deep imaging, including skin with a variety of pigments, bone surrounding the perimeter of the brain and over the midbrain-telencephalon boundary [28], curvature and pigmentation of the eye (particularly at telencephalic level), and muscle over the hindbrain region (Fig. 3).

**Figure 3:**
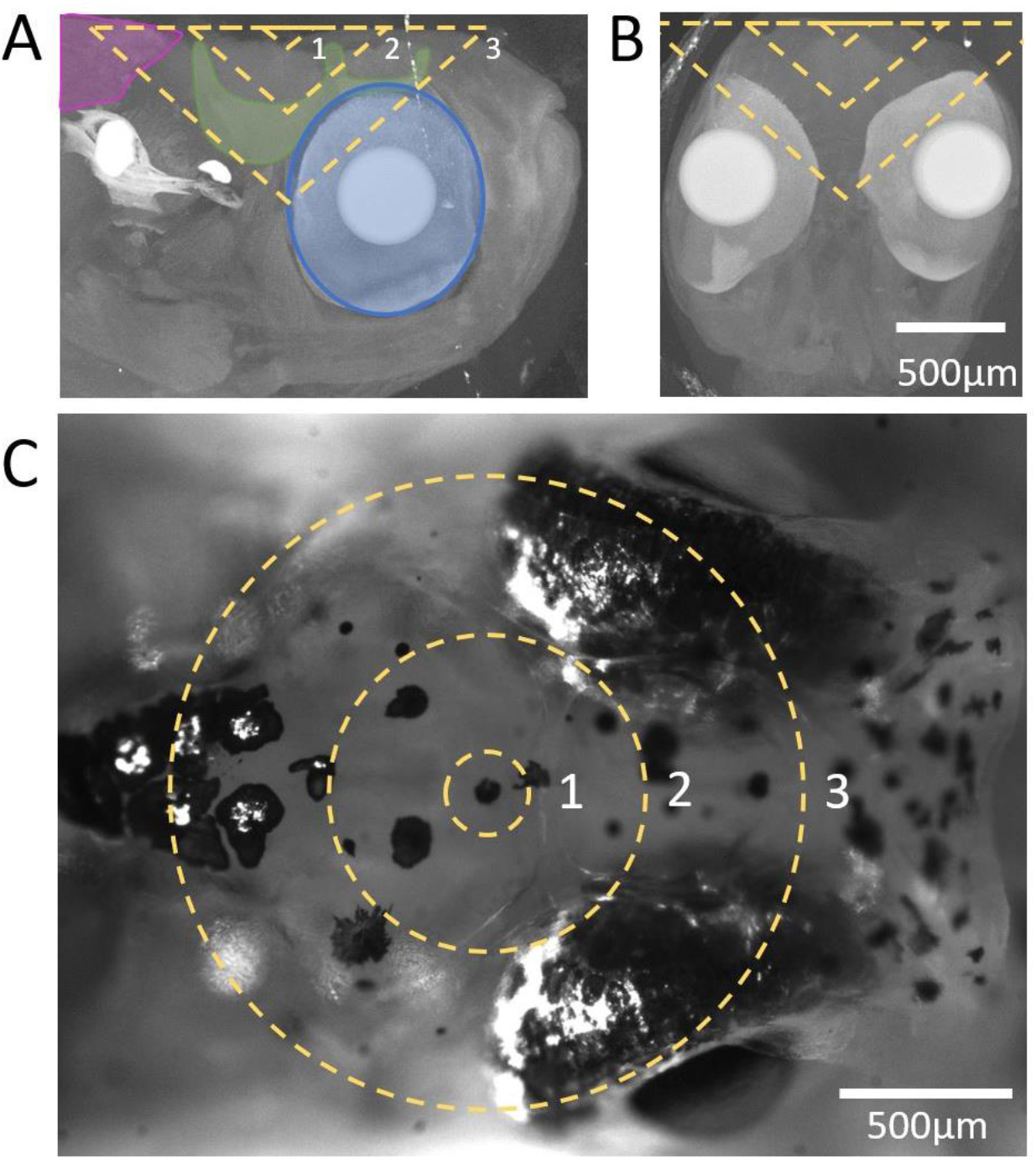
Demonstration of head structures that fall within the cone of light for deep imaging in the adult *D. dracula* brain. The dashed lines represent the cone of light for an NA of 1 at depths of 100 µm, 400 µm, and 800 µm labeled as 1, 2, and 3, respectively. A) Maximum projection of computed tomography (CT) images of the head that contain the right half of the brain. Muscle (magenta), bone (green), and eye pigments (blue) are outlined in the vicinity of the cone of light. B) Maximum projection of CT images of the head that contain the brain. C) White light image of the adult head.

In conclusion, we compare 2PM and 3PM performance for imaging through the entire depth of an adult vertebrate brain, in this case *D. dracula*. We show that higher order of excitation in 3PM not only increases contrast due to decreasing the excitation in out of focus regions, but also can help maintain a confined excitation focal volume by suppressing the side lobes of the PSF. The results further show that MPM also has the potential to further the application of optical imaging techniques to a wide range of aquatic vertebrates. Fishes alone account for more than half of all extant vertebrate species [29] and most do not currently have readily available genetic lines, but nonetheless are important for specific comparative questions regarding how brain diversity contributes to the evolution of behavior [30]. The small size of the adult *D. dracula* brain is particularly exciting for implementing whole-brain *in vivo* volumetric imaging with high temporal resolution. With MPM enabling optical access to adult brains and a repertoire of methods that allow observation of larval brains [9,31], aquatic vertebrates such as *Danionella* and other small species [19,20,32] become readily amenable for studying brain organization over the lifetime of a vertebrate for the first time.

## Methods

### Imaging setup

Images were obtained using a commercially available multiphoton microscope (Bergamo II system B242, Thorlabs Inc.) with a high numerical aperture objective lens (XLPLN25XWMP2, Olympus, NA 1.05). The back aperture was approximately backfilled for both 920 nm and 1280 nm excitation beams. The excitation source was a non-collinear optical parametric amplifier (NOPA, Spectra Physics) pumped by an amplifier (Spirit 1030-70, Spectra Physics). To control the excitation power, a half waveplate and a polarization beam splitter were used. A two-prism (SF11 glass) compressor was used to compensate for the normal dispersion of the optics of the light source and the microscope, including the objective lens. The full width at half maximum (FWHM) pulse width after the objective lens was measured to be 120 fs at 920 nm path and 50 fs at 1280 nm, assuming sech^2^ pulse intensity profile. For 3PM and 2PM, the excitation wavelength was 1280 nm and 920 nm, respectively. For deep imaging, low repetition rates of 333 kHz and 2 MHz were used for 3PM and 2PM, respectively. The FWHM pulse width was measured to be 120 fs at 920 nm path and 50 fs at 1280 nm, assuming sech^2^ pulse intensity profile. For some experiments a mode-locked Ti:Sapphire laser (Chameleon, Coherent) at 920 nm was used for 2PM excitation. The FWHM pulse width was measured to be 90 fs, assuming sech^2^ pulse intensity profile. Images were collected at approximately 1 frame per second over a field-of-view (FOV) of 539 µm by 539 µm with 512 by 512 pixels.

### Image Processing

Images were processed using ImageJ software. A 1-pixel radius median filter was applied to all images. For 3D reconstruction, the images were scaled by the ratio of z step size to the pixel size and were visualized using ‘Volume Viewer’ feature of the software. For line profiles, a 5-pixel wide line was drawn on the feature of interest. For resolution characterization line profiles were acquired using ImageJ and a gaussian fit was obtained using MATLAB.

### Characterization of Effective Attenuation Lengths (EALs)

For EAL characterization a 10-pixel wide line was drawn on blood vessels that had the top 1% of brightest pixel values. The brightest value of the line profile was used as the signal value for that depth. In areas where the autofluorescence of other features dominated the brightest pixel values, the image was cropped to only contain blood vessels and then the top 1% of brightest pixel values were chosen as the value for that depth. The signal value was then normalized to the square or cube of the power used at the corresponding depth, depending on the order of excitation.

### Characterization of Blood Vessel Distribution

Images were processed as described previously (see ‘Image Processing’). Areas corresponding to different regions (telencephalon, optic tectum, and cerebellum) were identified based on relative location in the brain and the distribution of vasculature. Areas of interest were outlined in ImageJ software and an appropriate threshold was applied to mask the blood vessels. Area fraction measurement function of ImageJ was used to obtain percentage of blood vessel for each frame, which likely over-estimates the blood volume concentration because the axial resolution (approximately 7 μm in our experiments) is comparable to or larger than the size of the capillary vessels. Therefore, the contribution to background fluorescence is less than the amount represented from images.

### Animals

Adult *D. dracula* were anesthetized in 0.025% benzocaine solution and their vasculature labeled via injection of a 10% solution of dextran fluorescein (70,000 molecular weight, ThermoFisher) and/or dextran Alexa Fluor 680 (70,000 molecular weight, ThermoFisher) into blood vessels in a highly vascularized region caudal to the operculum and rostral to the heart. Fish were stabilized by positioning them on a mountable putty (Loctite 1865809). Fish were perfused through the mouth and over the gills at a rate of 1 ml min-1 with an ESI MP2 Peristaltic Pump (Elemental Scientific) with well-oxygenated temperature-controlled fish system water containing 0.0185% benzocaine (4-l reservoir heated with a Top Fin Betta Aquarium Heater set to 25 °C). All procedures were in accord with the US National Institutes of Health guidelines for animal use in experiments and were approved by Cornell University’s Institutional Animal Care and Use Committee.

### CT (Computed Tomography)

Adult *D. dracula* were euthanized via deep anesthesia in 0.03% benzocaine solution. Whole fish were fixed in 4% paraformaldehyde at 4 °C for 24 h and then stained with 1% iodine metal and 2% potassium iodide. The stained fish was scanned at 120 kV per 10 W on the Zeiss Versa 520, using the ×4 objective and a resolution of 2.7 μm. The machine took 2,401 exposures of 0.7 s each, and the CT data was reconstructed using the standard Zeiss reconstruction software.

## Acknowledgements

Thank you to Joseph R. Fetcho for comments on the manuscript and Kristine E. Kolkman for helpful tips to improve experimental setup. We also thank the Cornell institute of Biotechnology for acquiring CT images of the adult *D. dracula* head.

