## Supplementary Figures for "Whole-brain optical access in small adult vertebrates with two- and three-photon microscopy"

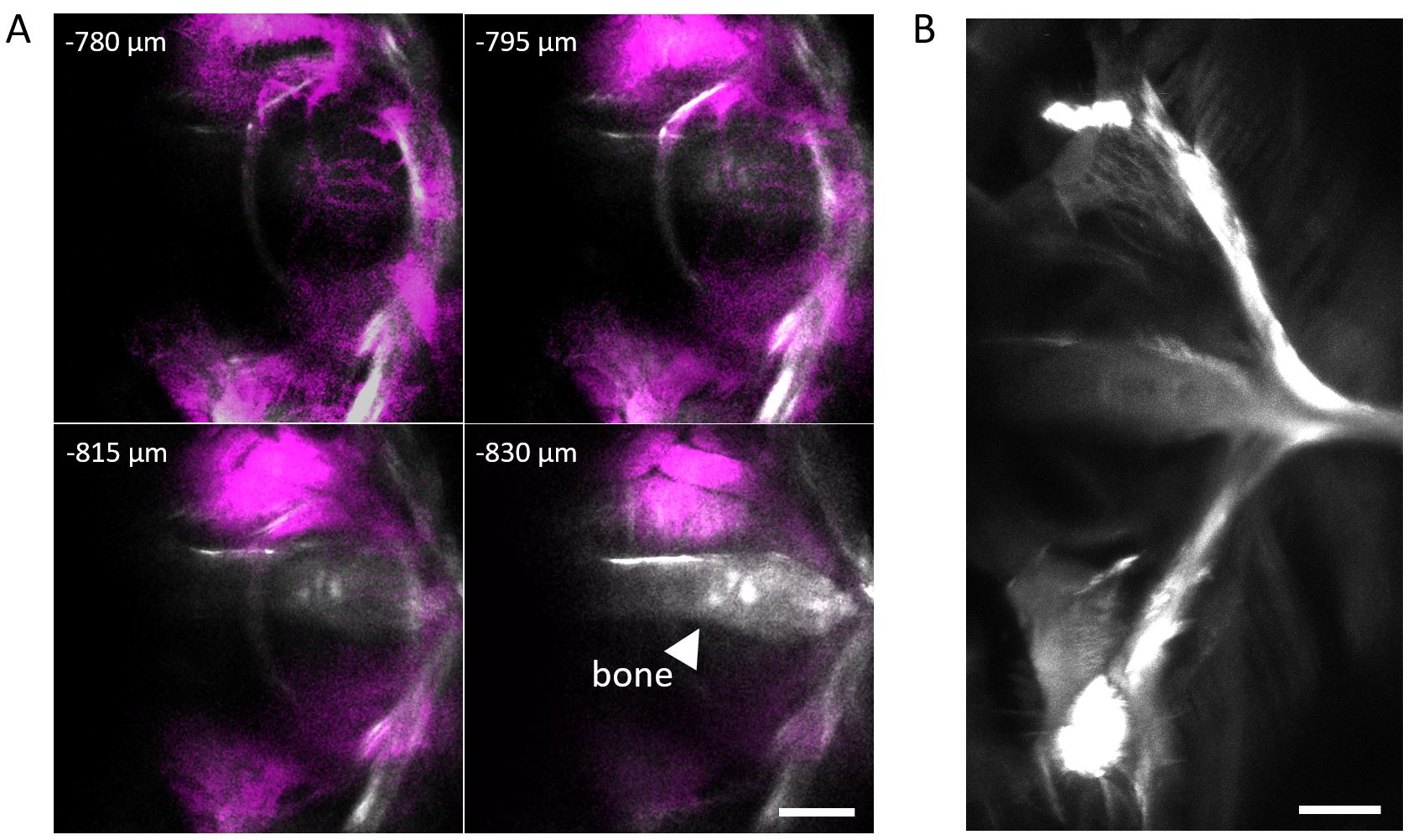

Supplementary Figure 1: Determining the bottom of the brain using SHG from the bone under the brain. A) Frames near the bottom of the brain containing three-photon excited fluorescence in magenta and SHG in white. The bone structure in the SHG channel is clearly visible at 830 µm depth as indicated by the arrowhead. The depth of the brain in this example was determined to be 795 µm which is the last image where clear blood vessel structure is visible. B) SHG of the entire bone (combination of two frames) under the brain. This structure matches previous characterization of *D*. *dracula* bone structure. Scale bars indicate 100 µm.

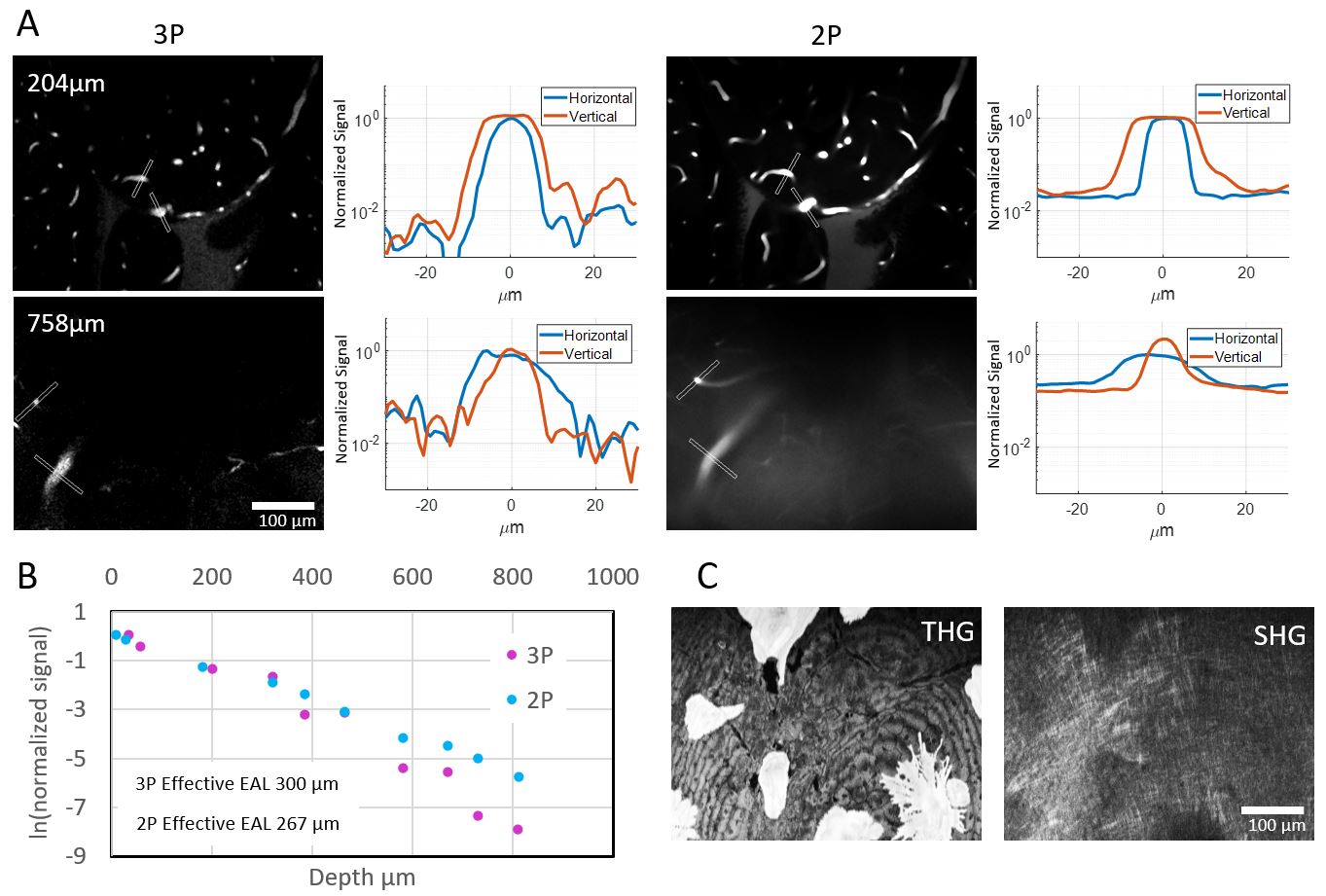

Supplementary Figure 2: Characterization of 2PM and 3PM images through the deepest part (midbrain) of the adult *D*. *dracula* brain. Excitation wavelengths of 920 nm at 80 MHz repetition rate and 1280 nm at 333 kHz repetition rate were used for 2PM and 3PM, respectively. To image the deepest part of the brain average power levels of 213 mW and 18 mW were used for 2PM and 3PM, respectively. A) Signal to background ratio comparison of horizontal and vertical blood vessels for 2PM and 3PM images at two depths inside the brain. In each line profile plot, the values are normalized to the maximum brightness of the horizontal blood vessel. B) Characterization of effective attenuation length inside the brain for 2PM and 3PM excitation wavelengths as described in the methods section. C) Maximum projection of THG (left) and SHG (right) of all frames containing the skin. Signals are generated by 1280 nm excitation. Pigments produce bright signal in the THG channel.

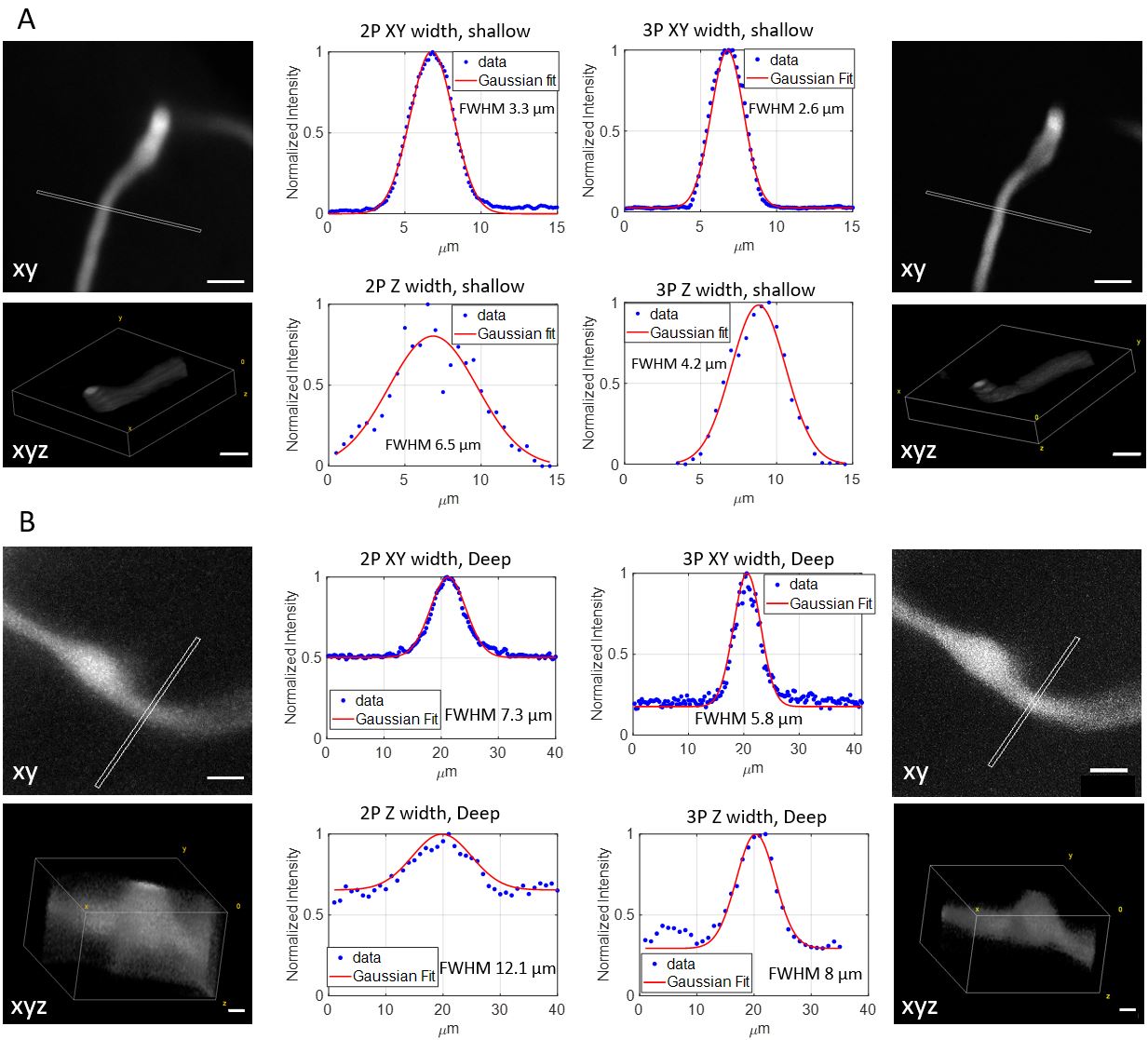

Supplementary Figure 3: Lateral and axial width measurements of a small blood vessel containing fluorescein (3PM, right panels) and Alexa Fluor 680 (2PM, left panels) in shallow (A) and deep (B) regions with 1280nm excitation.

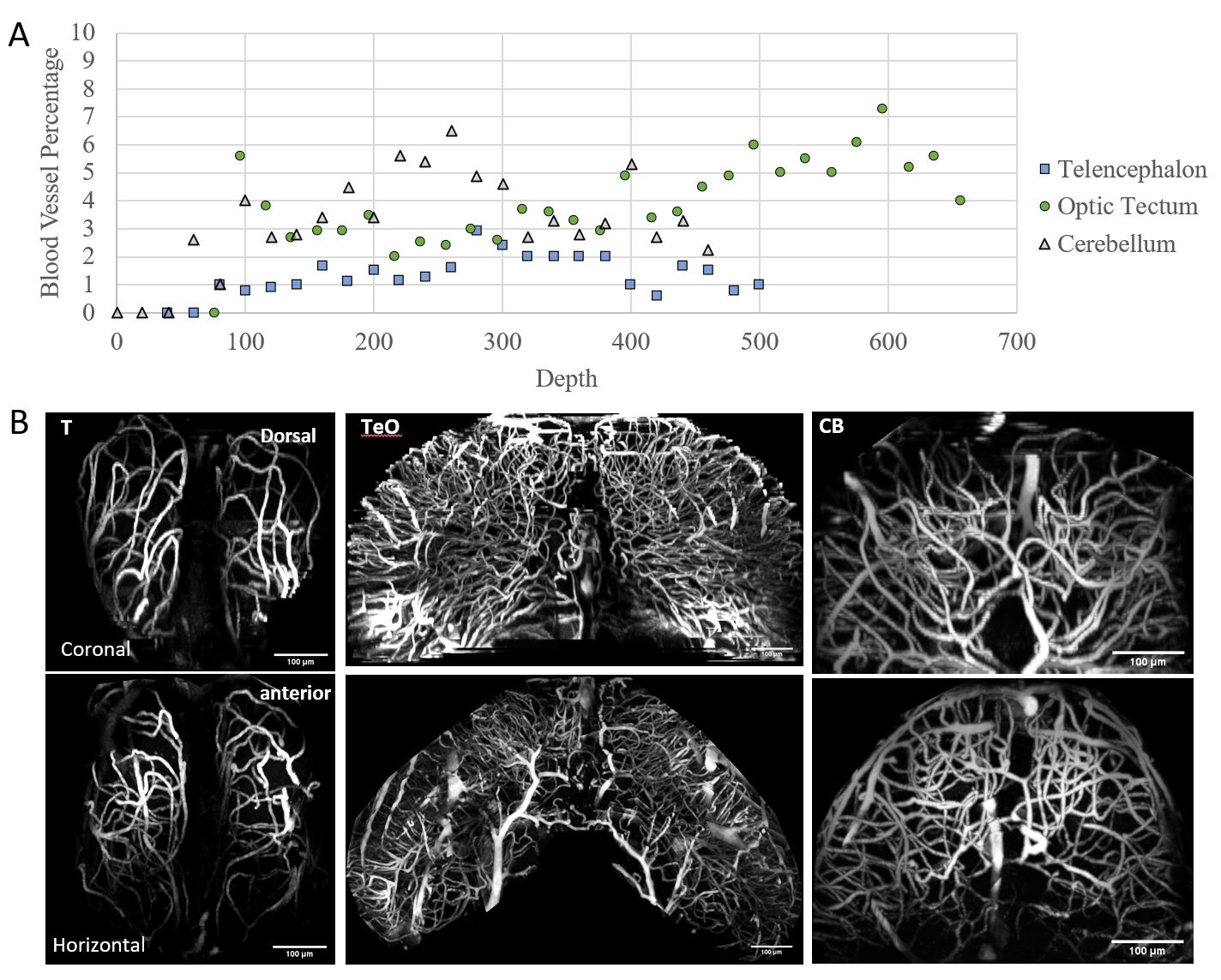

Supplementary Figure 4: Characterization of blood vessel density in different regions of the brain. A) Blood volume percentage in telencephalon, optic tectum, and cerebellum. characterized at various depths (see Methods). B) Maximum projection of vasculature in telencephalon (T), optic tectum (TeO), and cerebellum (CB). Coronal view of vasculature is demonstrated in top row. Horizontal view of vasculature is demonstrated in bottom row. Dorsal and anterior direction of each row is marked on the left-most image.

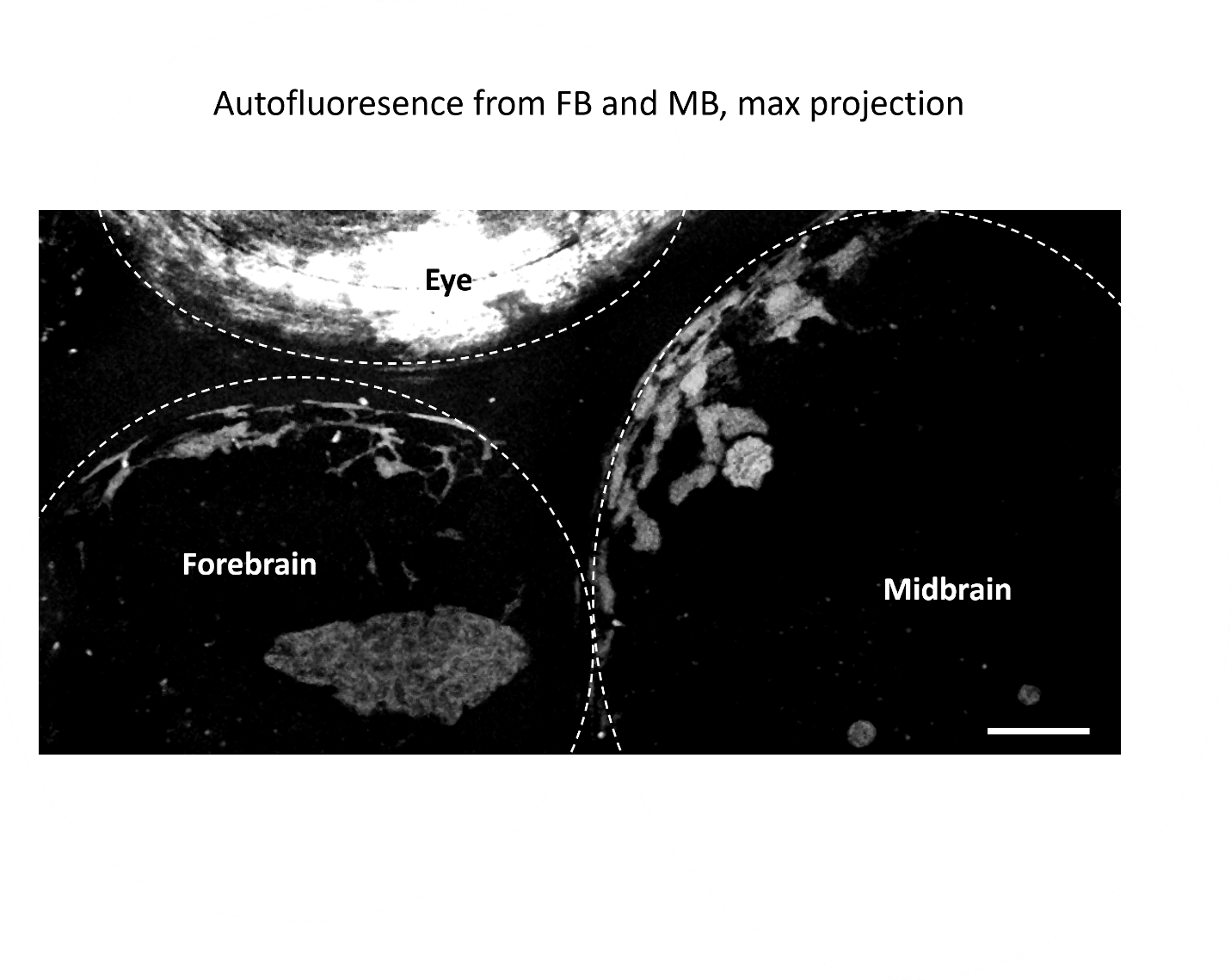

Supplementary Figure 5: Maximum projection of autofluorescence images collected through the brain. The eye, forebrain, and midbrain are outlined with dashed white lines. Images were collected without any dye injections administered to the fish. Scale bar indicates 100 µm.

 
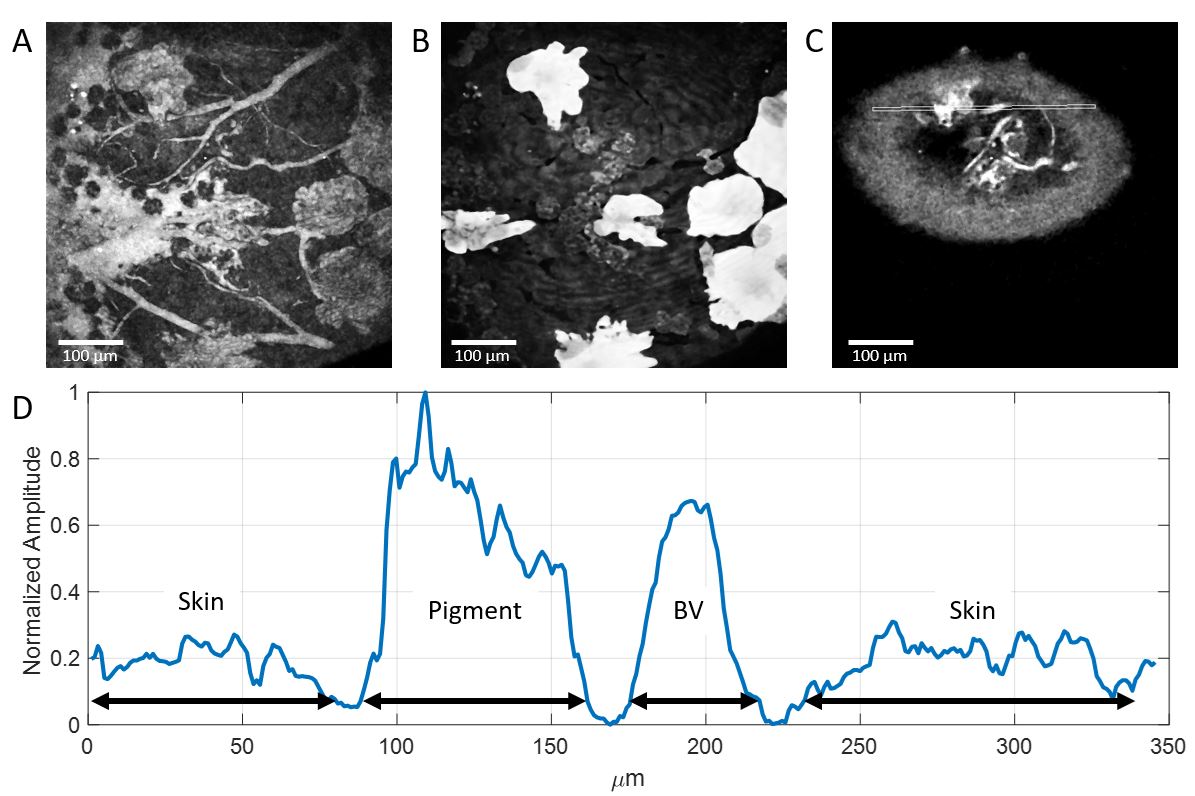

Supplementary Figure 6: Characterization of pigment and skin autofluorescence. All images are obtained with 1280 nm excitation. Vasculatures contain dextran-coupled Fluorescein. A) Maximum projection of all fluorescence images containing the skin. B) Maximum projection of all THG images containing the skin. C) A single frame containing fluorescence signal from skin (autofluorescence), pigment (autofluorescence), and blood vessels (BV, fluorescein). A 5-pixel-wide line is drawn over a region containing pigment, skin, and blood vessels. D) Normalized intensity of the line depicted in part C. Regions corresponding to pigment, skin, and blood vessel are marked with black arrowed lines.
